# Design and Preparation Of A Novel Hydrogel Based An Maleimide–Thiol Conjugation

**DOI:** 10.1101/2023.03.19.533362

**Authors:** Song Jiang, Monalisa Mohanty

**Affiliations:** Department of Biochemistry, Huzhou Institute Of Biological Products Co., Ltd. China; Environment and Sustainability Department, CSIR-Institute of Minerals and Materials Technology, Bhubaneswar – 751 013, Odisha, India

**Keywords:** Hydrogel, Maleimide–Thiol, Conjugation, Chitosan, Dextran

## Abstract

This article synthesized and characterized a novel hydrogel, which is formed using maleimide–thiol conjugation. Two precursors chitosan functionalized thiol groups and dextran functionalized maleimide groups prepared and characterized by NMR. The hydrogel studied by gelation time, swelling studies, viscoelastic properties, degradation rate. From gelation time result, we found that formed hydrogel gelation time could be changed with diffident weight percentage of precursors. Based on references, we found the best formular for the gelation and it was also determined for other studies. The swelling study indicated hydrogel has good flexibility and the degradation test indicated hydrogel is biodegradable. The viscoelastic test indicated hydrogel is elastic solid. From these studies, this a novel hydrogel could be potential for biomedical applications.

## 1. Introduction

Hydrogel is a water-swollen, crosslinked polymer network that is widely used in various fields, including medicine, agriculture, and engineering.[1] The hydrogel can be made from synthetic or natural polymers, such as polyethylene glycol, polyvinyl alcohol, or hyaluronic acid, among others.[2] One of the most significant advantages of hydrogel is its ability to absorb and retain large amounts of water. This property makes it an excellent material for wound healing, as it helps maintain a moist environment that facilitates faster healing.[3] Additionally, hydrogels have been shown to have excellent biocompatibility and low toxicity, making them an ideal option for use in various medical applications, including tissue engineering and drug delivery systems.[4] Hydrogels are also used in agriculture, where they can be used to improve crop yields and reduce water usage.[5] Hydrogel-based soil amendments can improve soil structure and water retention capacity, which helps plants grow in water-stressed environments. Additionally, hydrogels can be used to release fertilizers and pesticides in a controlled manner, reducing environmental contamination and increasing the effectiveness of these chemicals.[6] In engineering, hydrogels are used to develop smart materials that respond to environmental stimuli such as temperature, pH, or light. These materials have applications in areas such as sensors, actuators, and drug delivery systems.[7]

Hydrogel can be created through several methods, depending on the desired properties and applications of the final product. One of the most common methods is free-radical polymerization, which involves crosslinking monomers under the influence of free radicals. This process typically involves the use of a crosslinking agent, such as ethylene glycol dimethacrylate, and a free-radical initiator, such as ammonium persulfate.[8] Another method for creating hydrogel is through physical crosslinking, which involves using physical interactions, such as hydrogen bonding or electrostatic interactions, to form a polymer network. This method typically involves the use of natural polymers, such as chitosan or hyaluronic acid, and requires no chemical crosslinking agents or initiators.[9] A third method for creating hydrogel is through ionotropic gelation, which involves the use of ionic interactions to crosslink polymers. This method typically involves the use of a cationic polymer, such as chitosan, and an anionic crosslinking agent, such as tripolyphosphate, sulfobetaine.[10-14]

As we known, maleimide-thiol conjugation is a popular method for crosslinking hydrogels that involves the reaction of a maleimide functional group with a thiol functional group.[15-17] This method is widely used in the synthesis of hydrogels for biomedical applications, including drug delivery systems and tissue engineering.

Maleimide functional groups have a high affinity for thiol groups, making them an ideal choice for crosslinking hydrogels. The reaction between the maleimide and thiol groups forms a covalent bond, resulting in a stable crosslinked network. The reaction can be performed under mild conditions, which minimizes the risk of damage to bioactive molecules or cells encapsulated within the hydrogel.[18] One of the advantages of using maleimide-thiol conjugation to crosslink hydrogels is its high specificity. The reaction only occurs between maleimide and thiol groups, which reduces the likelihood of non-specific binding or unwanted reactions.[19] Another advantage of this method is its versatility. Maleimide functional groups can be incorporated into a variety of polymers, including synthetic and natural polymers, to create hydrogels with tailored properties. Additionally, thiol functional groups can be easily introduced into biomolecules, such as peptides or proteins, allowing for the creation of hydrogels with biologically active components.[20]

However, previous investigated hydrogels based on maleimide-thiol conjugation have many existing defects, including toxicity and slow gelation time. For instance, a typically example is from Zeng etc., who created a novel chitosan-PEG hydrogels from maleimide-thiol conjugation.[21] However, they utilized initiator UV lamp, which could be harmful to the cells in the biomedical application. And the gelation time at least 30 seconds with very high amount 20% precursors.

To conquer both defects mentioned above, in this paper, we selected two biocompatible nature materials chitosan and dextran. Then, we functionalized chitosan and dextran with shorter chain thiol groups and maleimide groups. Also, we didn’t use the initiator due to its toxicity. Meanwhile, we investigated conjugated hydrogel with its gelation speed, swelling behavior, viscoelastic property, and degradation rate.

## 2. Experimental Section

### 2.1. Materials

Glycine, maleic anhydride, glacial acetic acid, toluene, triethylamine, ethyl acetate, *N,N’-*dicyclohexylcarbodiimide (DCC), 4-dimethylaminopyridine (DMAP), p-toluenesulfonic acid monohydrate (PTSA) and anhydrous dimethyl sulfoxide (DMSO) were purchased from VWR. Magnesium sulfate, hydrochloric acid (HCl), ethanol and were purchased from Sigma-Aldrich. 4-(Dimethylamino)pyridinium 4-toluenesulfonate (DPTS) was synthesized from DMAP and PTSA. Deionized water was filtered via a Millipore purification device for all experiments.

### 2.2. Synthesis

#### 2.2.1 Synthesis of carboxyl end group maleimide (GMI)

**Scheme 1.**
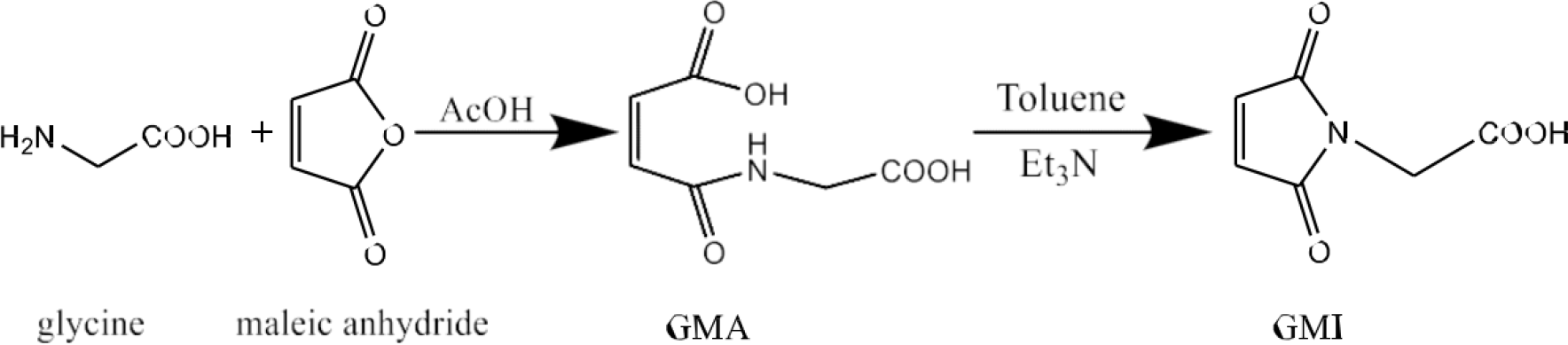
Synthetic route of GMI from maleic and glycine anhydride.

Scheme 1 outlines the procedure for preparing GMI. The method employed for synthesizing GMI was as described before. Typically, a mixture of maleic anhydride (0.1 mol, 9.8 g) in 50 mL of glacial acetic acid was added to a solution of glycine (0.1 mol, 7.5 g) in 100 mL of glacial acetic acid. The mixture was stirred at room temperature and then filtered. The resulting precipitate was washed with cold water and subsequently dried under vacuum. To a suspension of N-glycinyl maleamic acid (GMA) (16.8 mmol, 2.91 g) and Et_3_N (35.1 mmol, 3.55 g) in 500 mL of toluene, vigorous stirring was applied and refluxed for 1.5 hours. During this period, the byproduct water was removed through a Dean-Stark apparatus. The toluene solution was decanted away from the brown-colored oil and toluene was evaporated to obtain the triethylammonium salt. The resulting product was acidified with HCl to pH 2 and then extracted with ethyl acetate. The organic layer was dried with magnesium sulfate, and the ethyl acetate was subsequently evaporated to yield GMI.

#### 2.2.2 Synthesis of dextran with maleimide groups

**Scheme 2.**
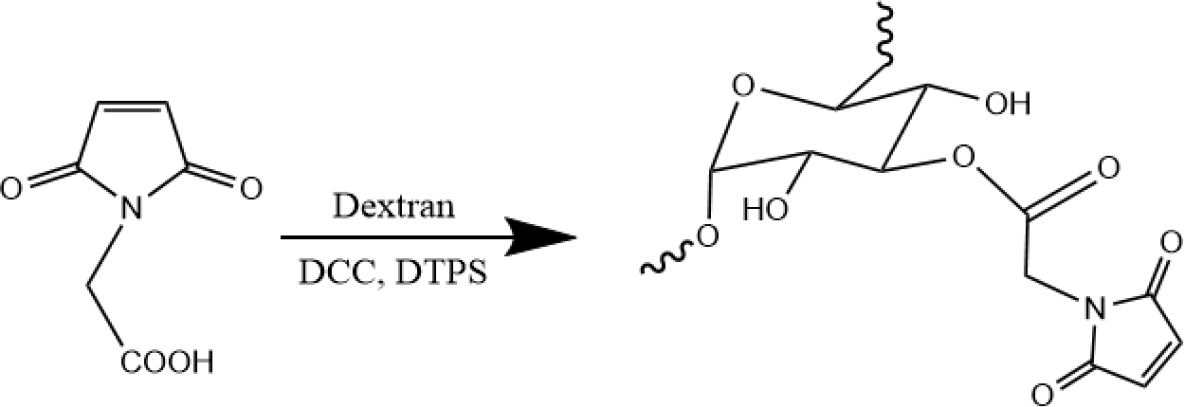
Schematic of synthetic route for dextran-MAL.

Scheme 2 outlines the process for synthesizing dextran (MW = 100 kDa) with maleimide groups (dextran-Mal). To create dextran-Mal, the hydroxyl group of dextran was esterified with N-maleoylamino acid using DCC mediation. The N-maleoylamino acid was prepared using a previously reported procedure. In this process, N-maleoylamino acids (0.72 g, 3.1 mmol) were dissolved in 20 mL DMSO, followed by the addition of DPTS (0.145 g, 0.45 mmol) and DCC (0.96 g, 4.65 mmol). Dextran (0.92 g, 5.15 mmol anhydroglucose (AHG) units) was dissolved in 10 mL DMSO and slowly added to the reaction mixture. After stirring at room temperature for 24 hours, the N,N’-dicyclohexylurea salt was removed by filtration, and the crude product was precipitated in cold ethanol. The precipitate was then collected by filtration, washed with ethanol, dissolved in water, and purified through ultrafiltration.

#### 2.2.3. Synthesis of chitosan with thiol groups

**Scheme 3.**
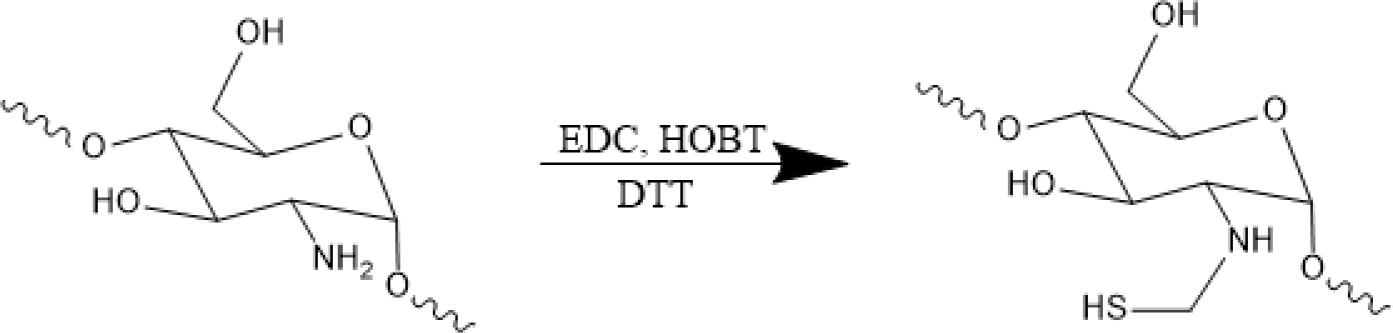
Schematic of synthetic route for chitosan-SH.

Chitosan with thiol groups (chitosan-SH) was produced and characterized using Scheme 3. To synthesize this compound, chitosan (MW = 100 kDa) was dissolved in DI water (5 mg/mL) and combined with three molar equivalents of cystamine dihydrochloride at pH 4.8. The conjugation process took place overnight after activation of the carboxyl group of chitosan using EDC and HOBT (at 3 equivalents) for two hours. The resulting product was purified using dialysis tubing with a molecular weight cutoff of 10,000 kDa against DI water for three days. To cleave the disulfide linkage of the cystamine component, the solution was treated with TCEP at 5 equivalents and stirred for two hours. The pH of the solution was then adjusted to 3.5 with HCl. Finally, chitosan-SH was precipitated in ethanol, re-dissolved in DI water, lyophilized, and stored at -20 ºC.

### 2.3. Hydrogel preparation

Dextran-Mal and chitosan-SH hydrogel precursors were dissolved in in pH 7.4 PBS, then followed by mixing dextran-Mal and chitosan-SH solutions. The maleimide groups “clicked” thiol groups to create the three-dimensional structure to form the gel.

### 2.4. Characterization

#### 2.4.1. Characterization of GMI, maleimide and thiol hydrogel precursors

Proton and carbon Nuclear Magnetic Resonance (1H and 13C NMR) were utilized to verify the molecular structures of GMI, maleimide, and thiol hydrogel precursors. The NMR spectra were recorded using an Avance Bruker with a BBO z-gradient probe. The experimental parameters included scanning each sample 128 times. The solvents employed were DSMO-d_6_ for GMI and D_2_O for dextran-Mal and chitosan-SH.

#### 2.4.2. Gelation time

The gelation time is determined using the vial tilting method. Briefly, dextran-Mal and chitosan-SH hydrogel precursors were dissolved in pH 7.4 PBS solution, respectively. The hydrogels were formed by mixing both solutions and vibrating quickly at room temperature. No flow within 1 min upon inverting the vial is regarded as the gel state.

#### 2.4.3. Swelling studies

Swelling studies of freeze-dried hydrogels were tested by a general gravimetric method. Samples were incubated at 37 °C in pH 7.4 PBS until weight is balanced. The swollen hydrogels were removed from the excess of water absorbed with a filter paper, and then weighed. The samples were then dried until constant weight. The swelling ratio was calculated using following Eq:

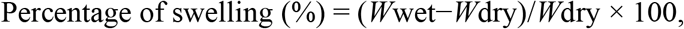

where *W*wet and *W*dry refer to the weight of wet and dry hydrogels respectively.

#### 2.4.4. Degradation

For degradation studies, weighted hydrogel freeze-dried samples were immersed in pH 7.4 PBS and incubated at 37 °C at selected time intervals. At each selected time interval, the samples were removed from PBS, dried and weighted. The hydrogel degradation was calculated using following Eq:

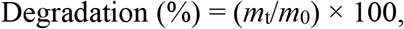

where *m*t and *m*0 refer to weight of freeze-dried hydrogels at selected time interval and before immersing in PBS respectively.

#### 2.4.5. Rheological analysis

Dynamic rheological behaviors of hydrogels are analyzed with a discovery hybrid rheometer, using parallel plate geometry (8 mm diameter). Frequency sweep measurements are performed at 37 °C from 0.1 to 100 rad s−1 at a fixed strain in the linear viscoelastic region.

## 3. Results and discussion

### 3.1. NMR Analysis

#### 3.1.1. GMI

GMI structure was confirmed by NMR spectra. In Figure 1. (a) ^1^H spectrum shows that the peaks around 13 and at 7.1 ppm are due to the -O-H bond of the carboxylic acid and the -CH=CH-bond of the maleimide ring.[2] In the Figure 2. (b) ^13^C spectrum illustrates that the peaks corresponding to carbonyl groups and the double bond of the maleimide ring appeared at 170.8 and 135.3 ppm, respectively. Furthermore, the peak of the carbonyl group of the carboxylic acid was detected at 169.3 ppm.

**Figure 1.**
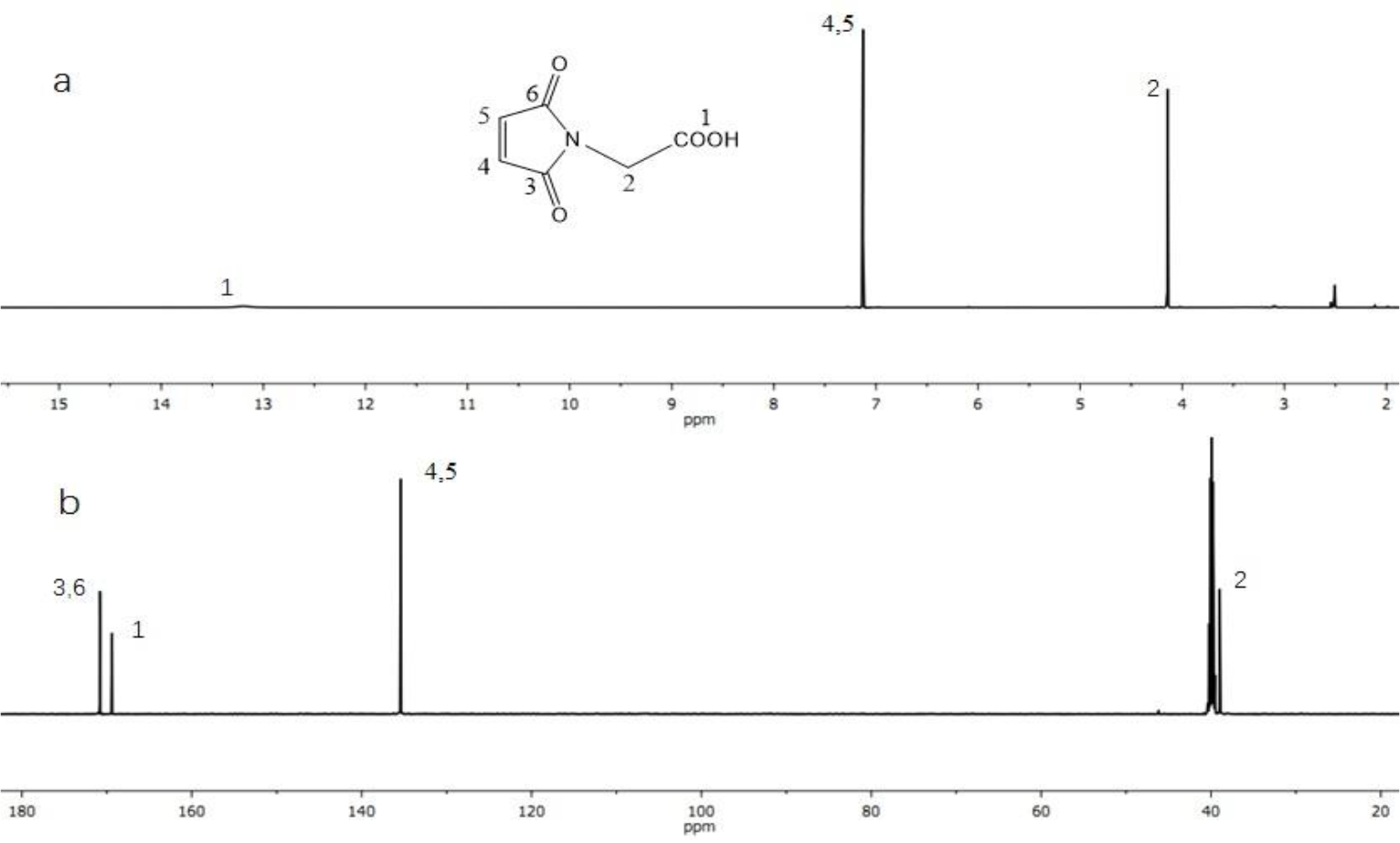
(a) ^1^H and (b) ^13^C NMR spectra of GMI in DMSO-d6.

**Figure 2.**
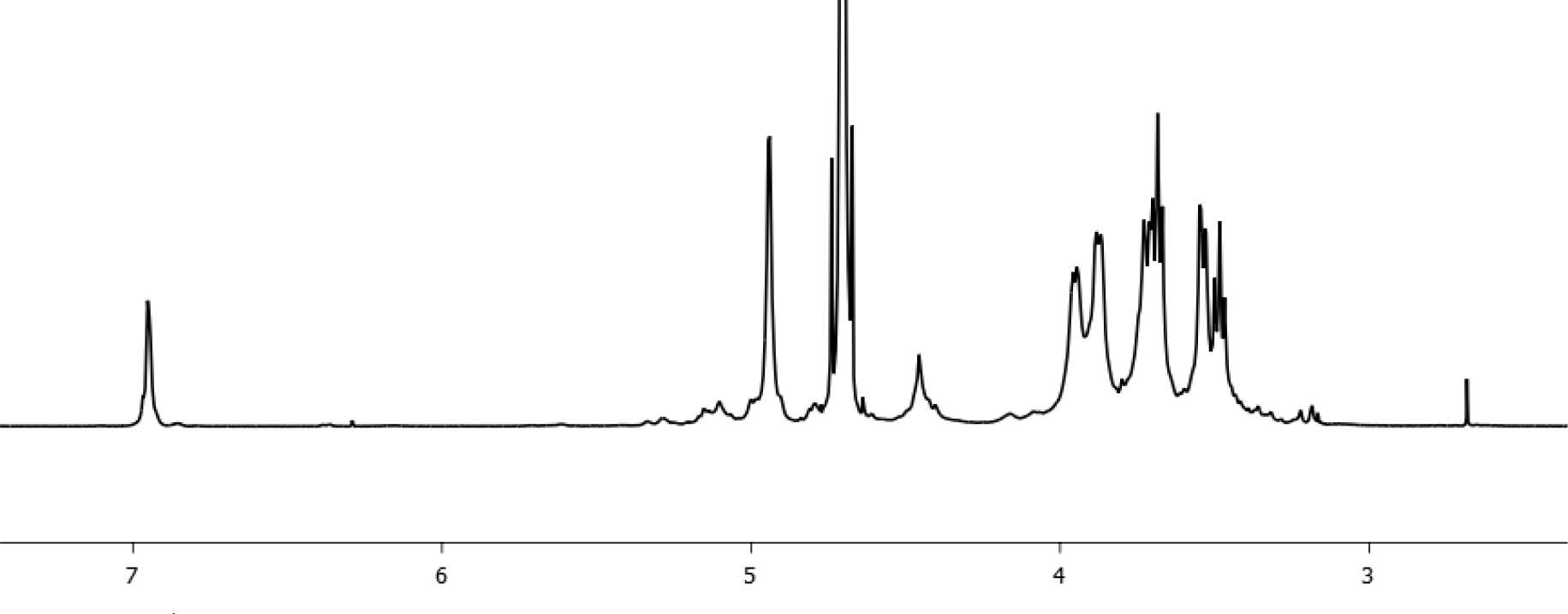
^1^H NMR spectrum of dextran-Mal.

#### 3.1.2. Dextran-Mal

The confirmation of the dextran-Mal structure has been established through the analysis of the ^1^H NMR spectrum. As illustrated in Figure 2, the peak observed at 6.9 ppm is indicative of the protons in the maleimide group, while the peaks detected at 4.0-3.4 ppm are attributed to the protons in the dextran backbone. Based on a comparison of the integrals of the resonance signals at 6.9 ppm and 4.0-3.4 ppm, the degree of functionality of the Dextran-Mal is estimated to be approximately 50%.

#### 3.1.3. Chitosan-SH

The confirmation of the chitosan-SH structure was achieved through ^1^H NMR analysis. The modification of chitosan was determined by utilizing the resonance of the acetamido group of chitosan as an internal standard at δ = 1.15 ppm. The peaks observed at δ = 1.95 ppm indicate the presence of cysteamine in chitosan-SH. Meanwhile, the peaks detected at 4.0-3.0 ppm are attributed to the protons in the chitosan backbone. Based on a comparison of the integrals of the resonance signals at 1.95 ppm and 4.0-3.0 ppm, the degree of functionality of the chitosan-SH is estimated to be approximately 70%.

### 3.2. Gelation time analysis

Table 1 summarizes the gelation time of hydrogels, with the weight percent of hydrogel precursors ranging from 1% to 2%. Gelation time ranges from ∞ (no gelation) to ≤1 second (ultrafast). The only feasible formula is 1.5% chitosan-SH and 1.5% Dextran-Mal because longer or shorter times could lead to the failure of cell encapsulation. For example, longer times may result in lower weight percent of hydrogel precursors used, leading to the formation of an amorphous gel. Conversely, shorter times could result in incomplete “clicking” of the maleimide and thiol groups, which may negatively impact cell growth. Consequently, this formula was used for the remaining experiments and analysis.

**Table 1.**
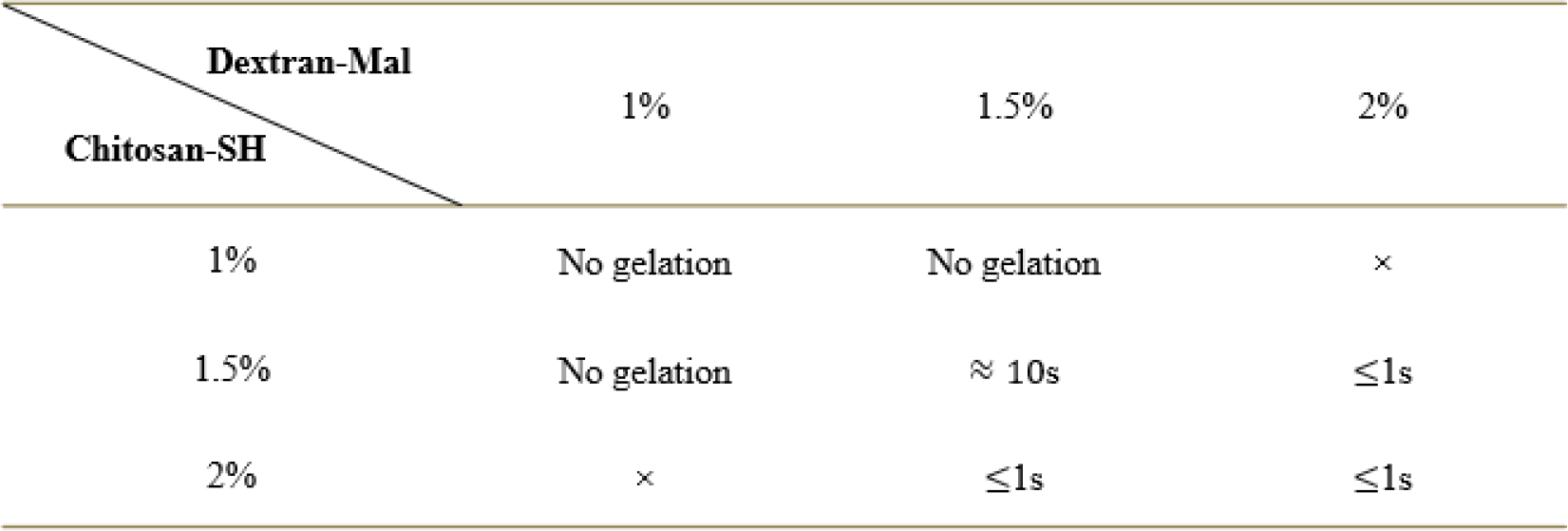
Summary of four hydrogels’ gelation time.

### 3.3. Degradation analysis

The weight loss of the hydrogels was plotted in figure 4. It exhibited that weight loss speed of hydrogels. The weight loss of hydrogels reached to 40% on the first week. After that day, the hydrogels were degraded with a steady rate until 23^rd^ day. Then, the hydrogels were degraded completely.

**Figure 3.**
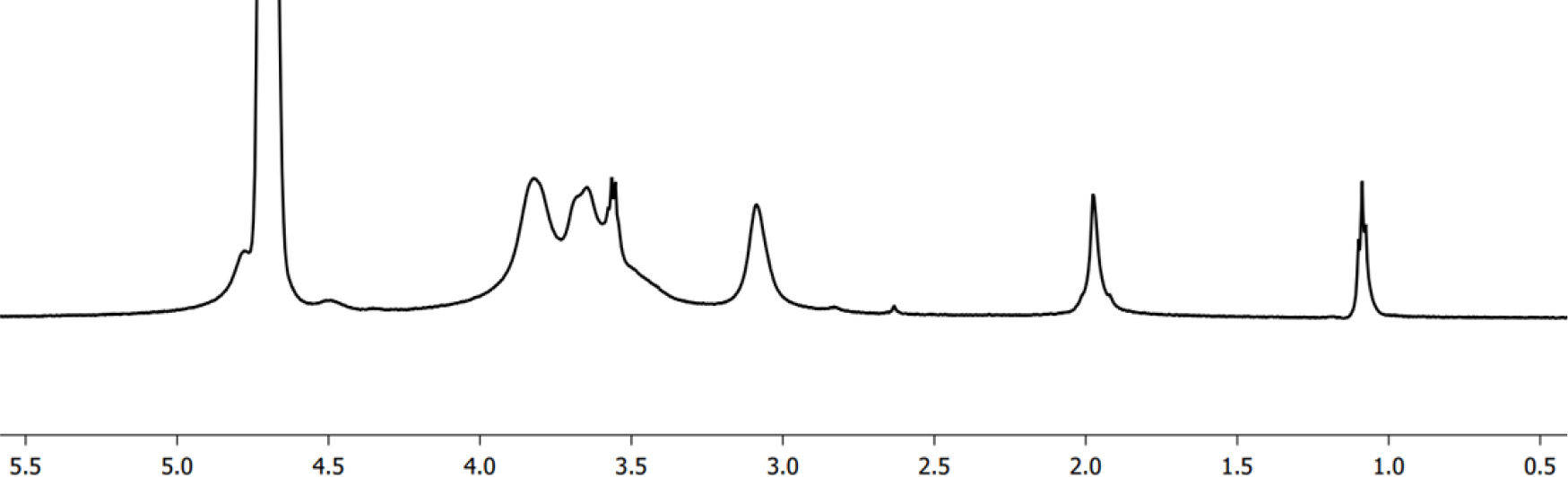
1H NMR spectra of chitosan-SH.

**Figure 4.**
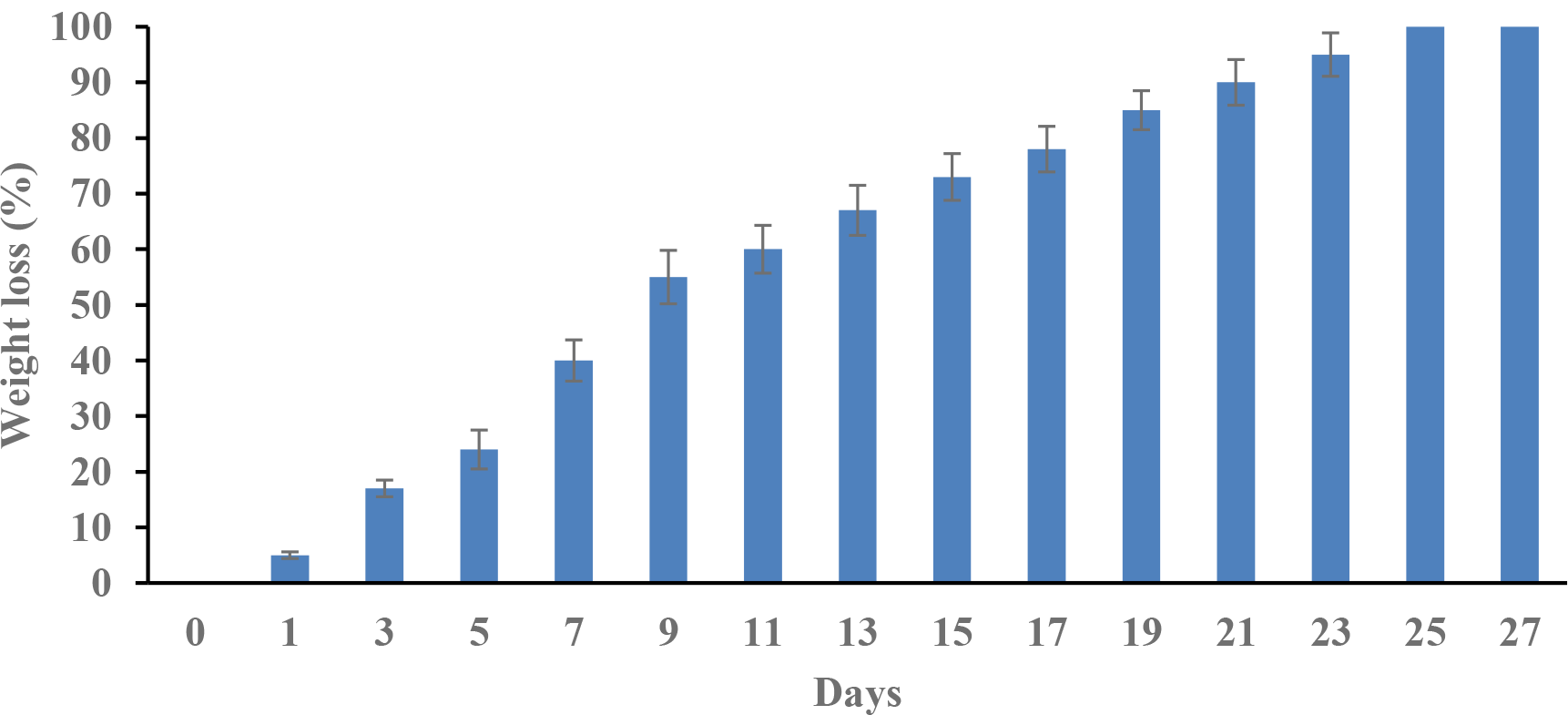
Weight loss of hydrogels at selected time intervals.

### 3.4. Swelling analysis

The swelling analysis is exhibited in Figure 5. The weights of freeze-dried hydrogels samples are from 1 gram to 5 grams. Since different types of hydrogels have different swelling properties. All samples of dextran-Mal and chitosan-SH hydrogels with different weights have similar swelling ratio, which is around 37%. The more hydrophilic samples will be submerged in PBS, the more water will be absorbed, and higher swelling ratio will be calculated. Here, it’s clear that the 1.5% dextran-Mal and 1.5% chitosan-SH hydrogels could absorb the water, which has the half mount weight as freeze-dried hydrogels. It indicates the hydrogels created by this formular have good flexibility ability to mimic extracellular matrix.

**Figure 5.**
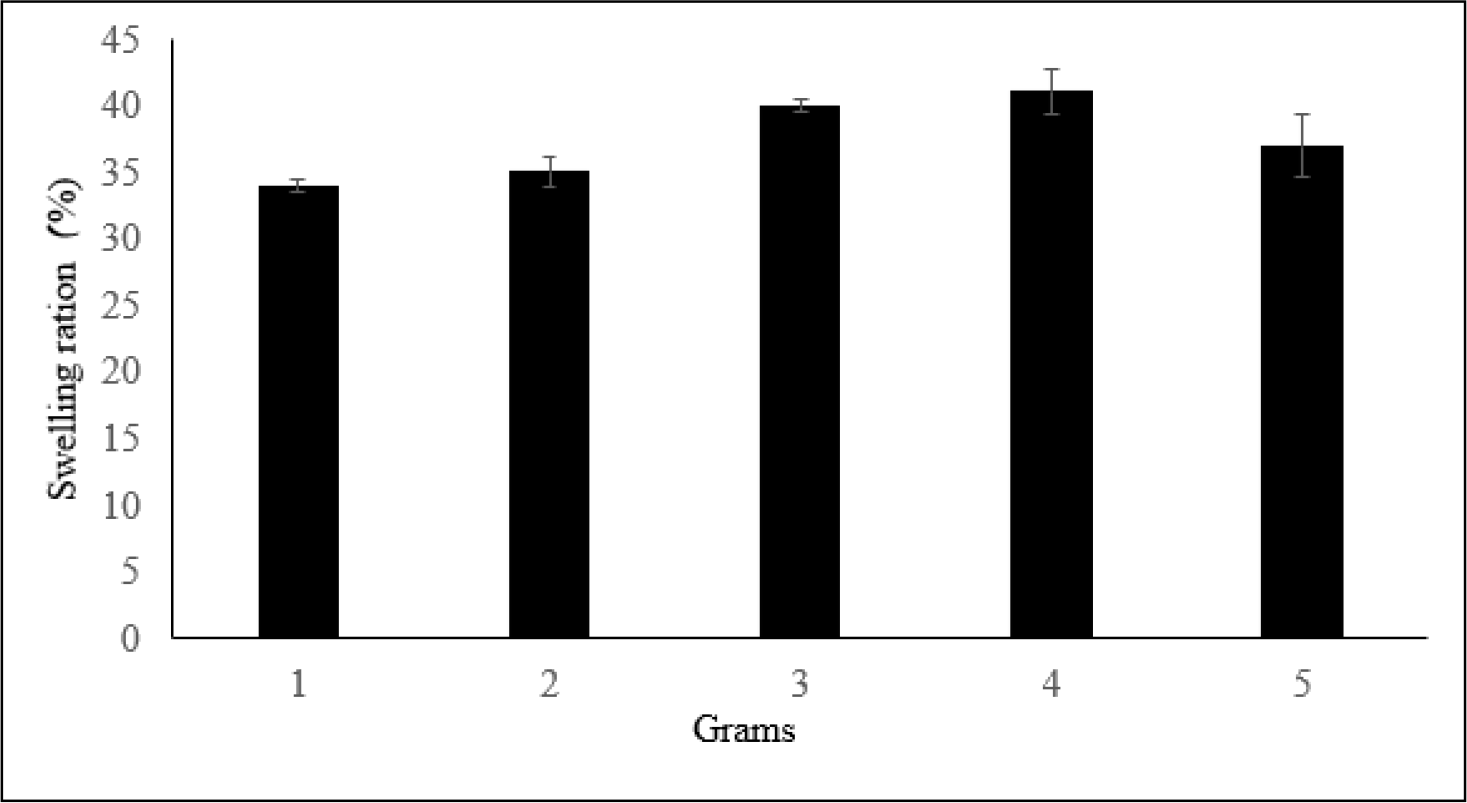
Summary of hydrogels’ swelling ratio.

### 3.5. Rheological analysis

**Figure 4.**
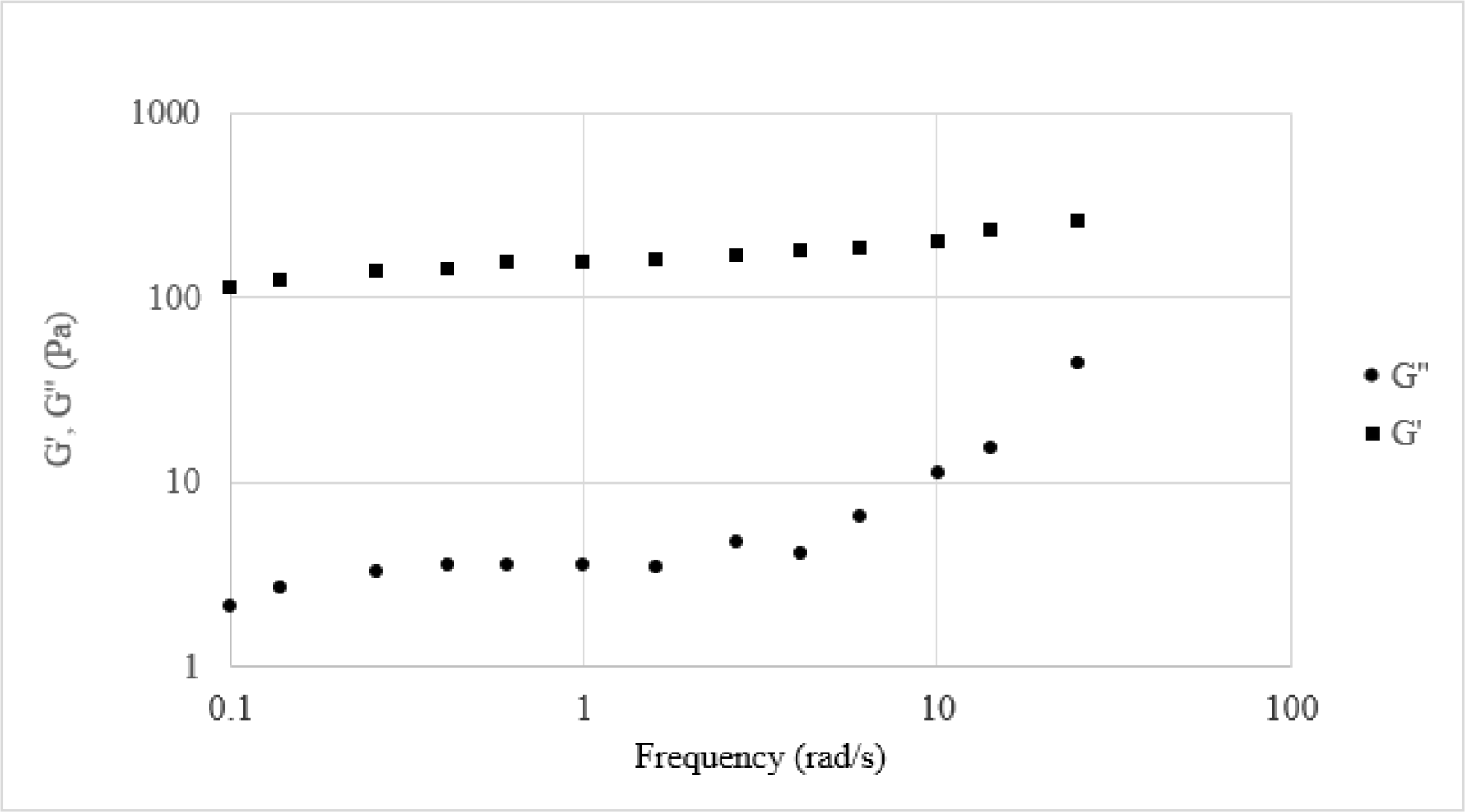
Frequency dependency of storage moduli (G′) and loss moduli (G′′) of the hydrogels.

The hydrogels were analyzed in Figure 4, with G′ and G′′ being plotted as a function of angular frequency. The results revealed that the hydrogels exhibited distinctive viscoelastic behaviors. Additionally, the moduli G′ were consistently higher than G′′, indicating that the hydrogels formed through maleimide-thiol coupling chemistry possess the properties of an elastic solid.

## 4. Conclusions

In this article, we conducted research to develop a novel biocompatible hydrogel by utilizing precursors functionalized with dextran-Mal and chitosan-SH through the Michael Addition. The successful synthesis of both precursors was confirmed by NMR analysis. The hydrogels were then subjected to a series of studies, which included the examination of their gelation time, swelling behavior, viscoelastic properties, and degradation rate. Based on our findings, we identified that the optimal formula was 1.5% dextran-Mal and 1.5% chitosan-SH, which we used for the remaining studies. Our findings suggest that the hydrogels possess good flexibility, as they were observed to mimic the extracellular matrix in the swelling study. Additionally, the hydrogels were found to completely degrade after 9 days in the degradation study. Moreover, the viscoelastic study indicated that the hydrogels formed via Michael Addition were elastic solids. Overall, our research provides evidence that this novel biocompatible hydrogel could have promising biomedical applications.

